# Applying Nanopore sequencing to a One-Health scenario for colistin resistance transmission among pigs, cows and the farmer

**DOI:** 10.1101/2019.12.20.884395

**Authors:** Joaquim Viñes, Anna Cuscó, Sebastian Napp, Judith Gonzalez, Ana Perez de Rozas, Olga Francino, Lourdes Migura-Garcia

## Abstract

One-Health studies applying massive-parallel and single-molecule sequencing are a suitable approximation to try to understand how antibiotic resistances flow between the human-animal-environment scenario. Colistin has been withdrawn in human medicine due to its toxicity, limiting its usage as a last-resort treatment option for multidrug-resistant Gram-negative bacteria. However, it is still used orally to treat *Enterobacteriaceae* infections in veterinary medicine. Since 2015, colistin resistance appeared to be located in mobile genetic elements, raising the concern of the likelihood of transmission by horizontal gene transfer between animals and humans. In this study, 202 faecal samples were collected in a mixed farm from pigs, calves, and the farmer. PCR for *the mcr-*1 gene was positive for 18 of the isolates, and Nanopore sequencing allowed us to determine the location of the gene, either on the chromosome or in plasmids. Three types of replicons were found within the positive isolates harbouring the *mcr-1*: IncX4, IncI2, and IncHI2. Four different genetic contexts probably indicate different stages of gene stabilization, either in the chromosome or plasmid, with *ISApl1* as the main insertion element flanking the gene. Moreover, 43 other resistance genes were found in our samples, related to more than six different antibiotic families (e.g. aminoglycosides, lincosamides, beta-lactams, macrolides, trimethoprim, phenicols, and sulphonamides). We found resistance genes against colistin and that six antibiotic families together in at least one of the isolates from human, swine, and bovine. Isolate 15B-22 harboured one plasmid with seven resistance genes related to four families of antibiotics other than polymyxins, meaning that there are more chances to maintain colistin resistance even with the withdrawn of colistin. Nanopore long reads allowed us to assemble the DNA elements within the isolates easily and determine the genetic context of the *mcr-1* gene. Furthermore, they allowed us to describe and locate more antimicrobial resistance genes to other antibiotic families and antiseptic compounds.

## Introduction

Single-molecule sequencing, such as Oxford Nanopore Technology (ONT), is gaining ground since its appearance into the market. The main characteristic from this type of sequencers is their ability to retrieve long reads, overcoming the problem of massive parallel sequencers and their short-read output. MinION™ is a portable and affordable sequencer that provides real-time data and results.

Long-read sequencing using ONT sequencers has been applied using different approaches to study the genetic content of bacterial samples, such as full-length 16S rRNA gene^1^, rrn operon sequencing^2–4^, metagenomics^5–10^ or Whole Genome Sequencing (WGS)^11,12^. Long-read output can span repetitive regions either from bacteria, archaea or eukarya, thus facilitating the assembly of genomes and the characterization of plasmids, and the location and genomic context of resistance genes^13^. These strategies have been also applied for resistome and plasmidome characterization^14–18^ since antimicrobial resistance has become a global concern.

Antimicrobials have been used globally in veterinary medicine for many decades. By means of natural selection, the consumption of antimicrobial agents has increased the selection of antimicrobial resistant bacteria in both human and veterinary medicine. Additionally, the presence of resistance genes in mobile genetic elements has probably played also a major role in the transmission of resistance inter- and intra-species.

The vast majority of the antimicrobials used in veterinary medicine are also used in human medicine. Hence, owing to its toxicity when applied systemically, colistin was withdrawn from human medicine. Nowadays, despite its toxicity, the emergence of multidrug resistant Gram-negative bacteria in hospital settings has left no other choice but to use colistin as the last-line treatment option. Contrarily, in veterinary medicine, colistin sulphate has been used orally for many decades to treat infections caused by *Enterobacteriaceae*^19^. In particular, colistin tablets are available for calves for the prevention and treatment of neonatal colibacillosis^20^. Additionally, studies performed in different EU countries have reported the prophylactic and metaphylactic use of colistin for the prevention and treatment of enteric diarrhoeas of pigs^21–24^. In this sense, Spain was the country with the highest sales of colistin for food producing animals in the EU in the year 2014^25^. Fortunately, consumption of colistin has been drastically withdrawn in the last years after implementation of specific programme targeting pig production and the voluntary agreement of producers^26^.

Until 2015, resistance to colistin was associated to mutations in the chromosome leading to changes in the charge of the lipopolysaccharide that modify the target and reduce the binding of the antimicrobial^27^. More recently, different plasmid mediated mechanisms conferring resistance to colistin have been identified^28–32^, with *mcr*-1 distributed worldwide^33^. The emergence of colistin resistance located in mobile genetic elements has raised the concern of the scientific community since transmission of resistance from farm to fork could amplify the difficulty of treating severe infections in human hospitals.

Back in 2017, co-occurrence of *mcr*-1 and *mcr*-3 was described for the first time in an *Escherichia coli* of calf origin^34^. This isolate was obtained from a faecal sample at slaughterhouse during the Spanish National Monitoring Programme back in 2015. The farm providing the calf was identified and a visit was carried out in September 2017 to determine if this *E. coli* genotype was endemic at the farm. In this context, we have applied nanopore sequencing from ONT to assess the suitability of this technology to study the transmission of colistin resistance with a One Health approach. Results obtained of this cross-sectional study are described in this manuscript.

## Material and Methods

### Study designed

After the identification in 2017 of an *E. coli* isolate of bovine origin harbouring both *mcr*-1 and *mcr*-3, the farm from where the sample was obtained was contacted to follow up on the finding. This farm belonged to an independent Spanish producer and contained two separate areas. The first area was a farrowed to finished system for pig production. Not more than 100-meter distance and without a physical barrier was a multi-origin bovine fattening farm also property of the farmer.

The number of samples to be collected in each of the housing facilities was calculated to allow the detection of a prevalence of *mcr-1-mcr-3-E. coli* of at least 5%, with a 95% confidence level. Sample size calculations were carried out using the WinEpi tool (http://www.winepi.net/uk/index.htm).

Additionally, the farmer was interviewed by the research team and the farm’s treatment book was inspected to determine the antimicrobial treatments prescribed to the two animal species from 2015 to the time of sampling.

### Isolation of colistin resistant E. coli and mcr-detection

Faecal samples were taken from individual animals and transported to the laboratory at 4° C on the day of sampling. Faeces were homogenized and plated onto both, McConkey agar and McConkey agar supplemented with colistin (2mg/L). For quality control of the colistin plates, a positive and negative control were also included. Following incubation, one presumptive *E. coli* isolate per positive sample was identified by PCR^35^ and stored at −80° C for further analyses. Detection of five different *mcr* genes was performed by multiplex PCR as described before^36^.

### Antimicrobial susceptibility testing

Minimal inhibitory concentration was carried out for 14 antimicrobial agents (VetMIC GN-mo, Swedish National Veterinary Institute) in those isolates harbouring *mcr*-genes. Antimicrobials tested were ampicillin (1 to 128 mg/L), cefotaxime (0.016 to 2 mg/L), ceftazidime (0.25 to 16 mg/L), nalidixic acid (1 to 128 mg/L), ciprofloxacin (0.008 to 1 mg/L), gentamicin (0.12 to 16 mg/L), streptomycin (2 to 256 mg/L), kanamycin (8 to 16 mg/L), chloramphenicol (2 to 64 mg/L), florfenicol (4 to 32 mg/L), trimethoprim (1 to 128 mg/L), sulfamethoxazole (8 to 1,024 mg/L), tetracycline (1 to 128 mg/L), and colistin (0.5 to 4 mg/L). Epidemiological cut-off values were those recommended by the European Committee on Antimicrobial Susceptibility Testing (EUCAST). Multiresistant was defined as resistance to at least three different antimicrobial families^37^.

### DNA extraction

DNA from *mcr*-positive isolates was extracted using QIAGEN DNeasy^®^ Ultraclean Microbial Kit under manufacturer’s conditions.

### Nanopore sequencing library and data preparation

DNA was quantified using Qubit dsDNA BR assay (Invitrogen by ThermoFisher Scientific). Sequencing was performed using MinION™ sequencer from Oxford Nanopore Technologies (ONT) including two runs with 9 barcoded isolates in each. Two sequencing libraries were prepared using 400 ng DNA per sample with the Rapid Barcoding Kit (SQK-RBK004) following the ONT protocol.

The samples were run using MinKNOWN software (version 18.07.18). Fast5 files generated were basecalled and demultiplexed (sorted by barcode) using Albacore v2.3.1 to obtain fastq files. Reads classified as pass (minimum Phred score of 7) were used for further steps. A second round of demultiplexing was performed with Porechop^38^ in which barcodes that agreed with Albacore were kept and the rest removed. Moreover, Porechop was used to trim barcodes, chimeric reads and sequencing adapters from the sequences.

### Assemblies

Once reads were pre-processed, Unicycler^39^ was used to assembly the genomes. Unicycler allows working with only Illumina reads, only Nanopore reads or in a hybrid way merging both types of reads. We applied Unicycler using long reads, which assembles using miniasm^40^ and minimap2^41^ and polishes with Racon^42^.

### Data analysis

Contigs were annotated using Prokka^43^ and visualized using Bandage^44^. NCBI BLAST tool was used within Bandage for visualizing specific genes. Mauve^45^ was used for visualizing the alignment of the contigs. Abricate^46^ along with different databases (CARD, NCBI and PlasmidFinder) was applied for antibiotic resistance gene and plasmid replicon description. Resistance genes with at least 90% of both coverage and identity were kept.

## Results

Occurrence of colistin resistant E. coli

A total of 202 faecal samples were collected from pigs (n=57), calves (n=144) and the farmer (n=1). Of them, 41 presented growth on the McConkey supplemented with colistin and were confirmed to be *E. coli*. Multiplex PCR for detection of the five *mcr*-genes confirmed the presence of 18 samples positive for *mcr*-1 gene. Of those, 13 were isolated from calves, 4 from pigs and 1 from the farmer. No other *mcr* genes were detected.

### Antimicrobial susceptibility testing

As can be seen in Supplementary Table 1, MIC values for colistin varied with one isolate exhibiting a MIC of 2 mg/L, 15 isolates equal to 4 mg/L and 2 showing a MIC ≥ 8 mg/L (Supplementary Table 1). Furthermore, 100% of isolates were resistant to ampicillin, ciprofloxacin, streptomycin, chloramphenicol, sulphametoxazole and trimethoprim. Additionally, 95% exhibited resistance to tetracycline, 89% to nalidixic acid and florfenicol, 78% to kanamycin and 72% to gentamicin. Finally, phenotypic resistance to cefotaxime was observed in three isolates whereas resistance to ceftazidime was detected in two. All isolates were multiresistant.

### mcr-1 harbouring contigs, gene location and replicons

Sequencing results regarding colistin resistance were in agreement with phenotype and PCR results, except for one isolate (15A-11) where *mcr*-1 could not be detected (Table 1). Two of the isolates (P2-16 and 15B-13) carried the *mcr*-1 gene inserted in the genome without the presence of any replicons around the region where the gene was located (Table 1). These isolates were from swine and bovine origin, respectively. The remaining 15 isolates have the *mcr*-1 gene located in a contig carrying replicons: IncX4, IncI2 and IncH2. The most abundant replicon associated with *mcr-1* gene was IncX4 (n=13), detected in isolates from pigs, calves and the farmer. One isolate of bovine origin (15A-1) carried *mcr*-1 associated with IncI2. Finally, another isolate of bovine origin (15B-22) harboured two copies of the *mcr*-1 gene associated with IncHI2 and IncX4 replicons, respectively. Plasmid sizes ranged from 33,196 for IncX4 (mean 34,700 bp) up to 61,594 bp for IncI2 and 233,608 pb for IncHI2.

**Table 1.**
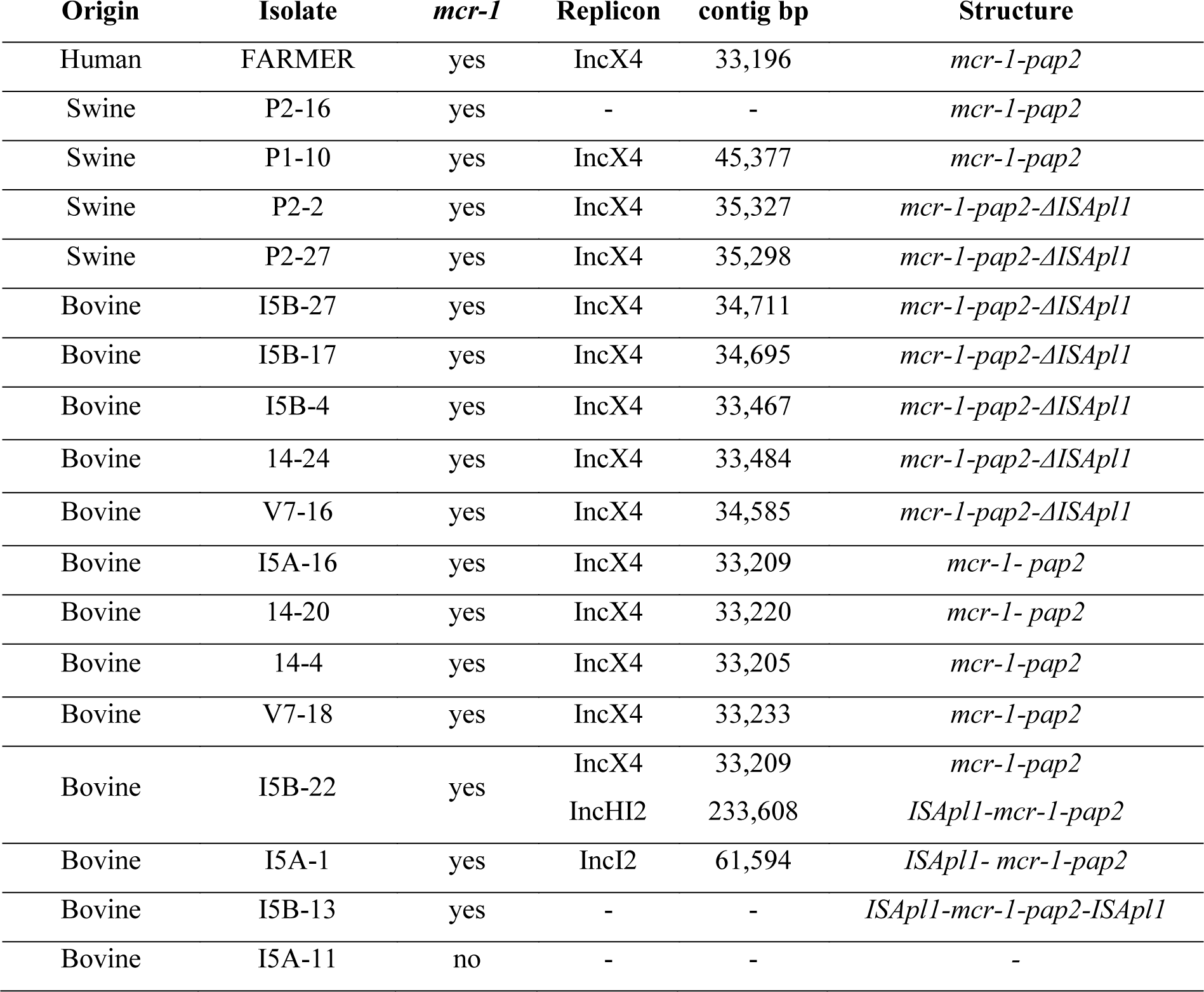
Identification of *mcr-1* gene, its genomic structure, and the associated replicons in the farm isolates from bovine, porcine or human origin.

### Genomic context of mcr-1

The genomic context of the *mcr*-1 gene was compared between the different isolates. Four different configurations of elements *ISApl1, pap2, mcr1* and *ΔISApl1* were detected (Table 1 and Figure 1). The most common structure (n=8) contained the *mcr-1* gene upstream the *pap2* with no signs of ISApl1 upstream or downstream. This structure was always integrated in IncX4 replicons or in the genome, and was detected in isolates of bovine, swine and human origin.

**Figure 1.**
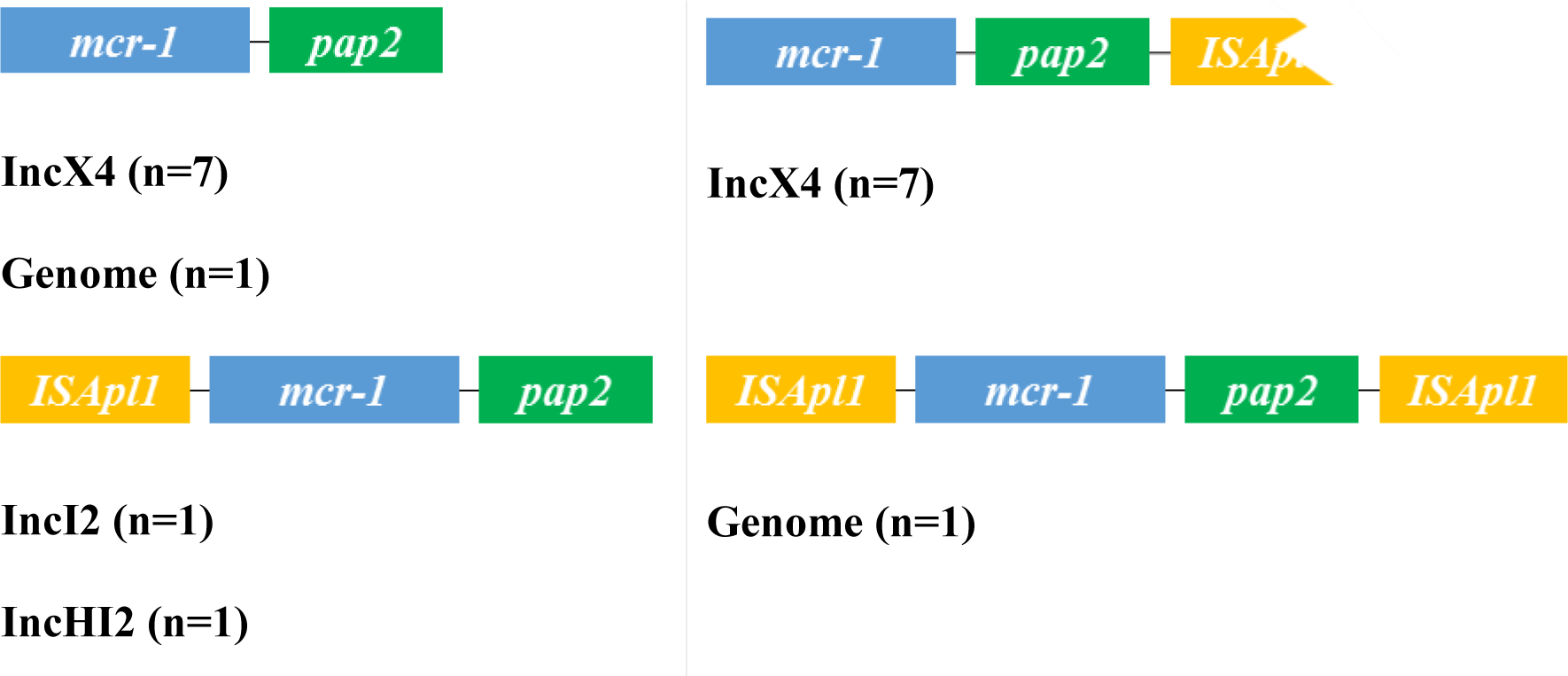
Genomic context of the *mcr*-1 region with 4 different configurations of elements *ISApl1, pap2, mcr1 and ΔISApl1*. In yellow, full or fragmented ISApl1 element; in blue, *mcr-1*; in green, *pap2*.

The second more frequent combination (n=7) also associated to IncX4 plasmids contained *mcr-1, pap2* and a small fragment of ISApl1 element (218 bp) downstream *pap2*. Two other isolates contained *mcr*-1 flanked by ISApl1 (upstream) and *pap2* (downstream), respectively. In these cases, the replicons associated to the *mcr-1* genes were of IncI2 and IncHI2 families. Finally, one isolate (15B-13) presented ISApl1 flanking the *mcr-1-pap2* construction. This turned out to be an isolate of bovine origin in which the construction was located in the genome.

### Other replicons and antibiotic resistance genes retrieved by Nanopore sequencing

In total, twenty-two replicons were identified and are shown in Figure 2. Each one of the replicons was associated with a single plasmid, with the exception of lncHI2 and lncHI2A that were found in the same plasmid in all the isolates. Twenty-one out of the twenty-two replicons were detected in bovine isolates; five in pig isolates and seven in the farmer. The number of replicons per isolate ranges from two (pig isolates) to nine (bovine isolate 15B-13 with *mcr-1* gene located in the genome).

**Figure 2.**
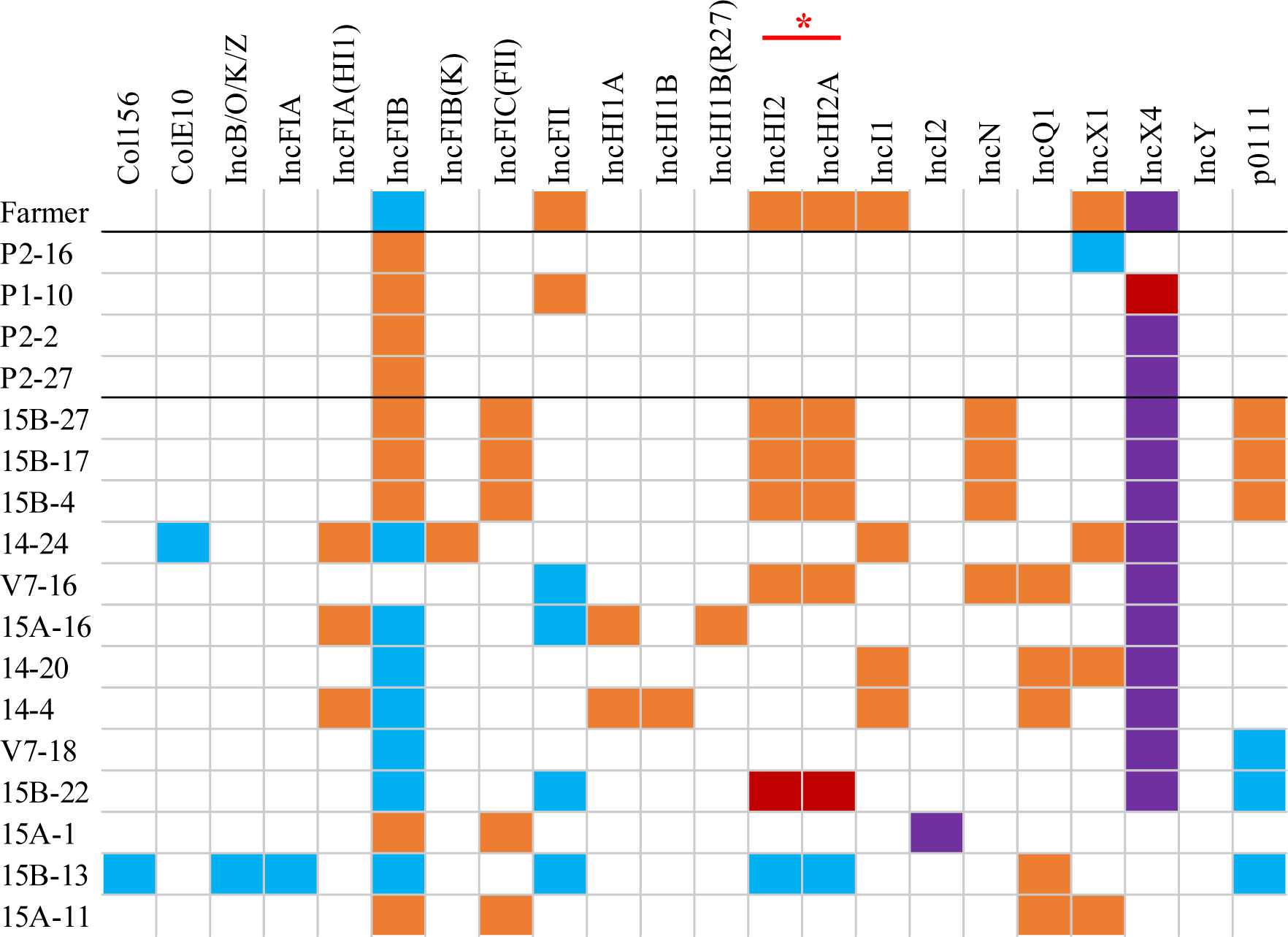
Replicons found in each isolate: contigs harbouring *mcr-1* gene (**violet**); *mcr-1* and other antimicrobial resistance genes (**red**); other antimicrobial resistance genes instead of *mcr-1* (**orange**), and contigs that do not present any antimicrobial resistance gene (**blue**). *mcr-1* gene is located in the genome in isolates 15B-13 and P2-16. *For each strain that presents both replicons, these are found in the same contig.

Table 2 shows all the genes retrieved applying Abricate on Nanopore contigs: 100% of the samples present genes conferring resistance to beta-lactams, aminoglycosides, phenicols and tetracyclines. 94.4% of the isolates harboured genes conferring resistance to sulphonamides, and 83.3% to trimethoprim. 38.9% of the isolates present genes against macrolides and lincosamides.

**Table 2.**
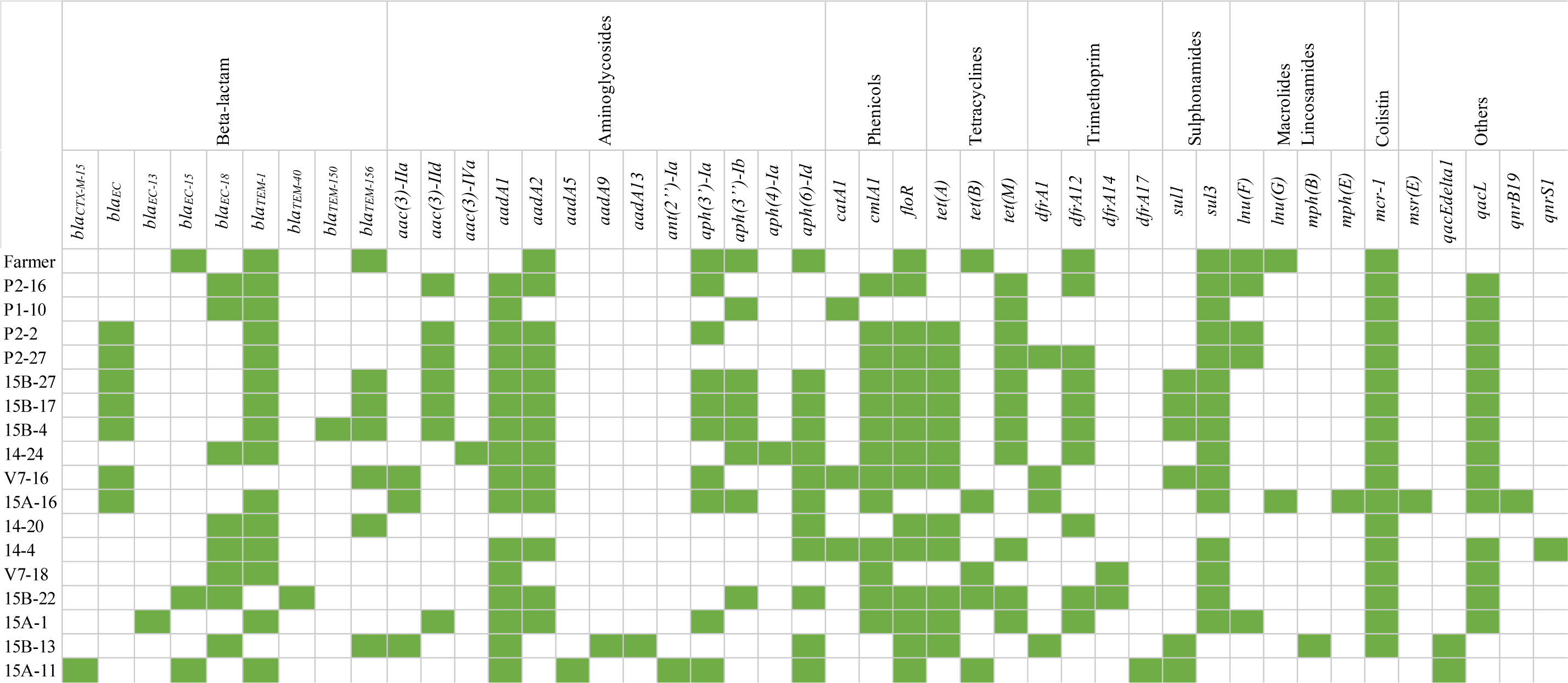
All genes retrieved with abricate from all isolates.

Besides *mcr-1*, two of the plasmids harboured other resistance genes (Figure 3). P1-10 isolate also presented a *tet*(M) gene, which conferred resistance to tetracycline. Whereas the mean length of contigs harbouring IncX4 was 34,700 bp, this contig was 45,377 bp, presenting an inserted region where *tet*(M) laid. On the other hand, as mentioned above, bovine isolate 15B-22 presented two plasmids with different replicons associated to *mcr-1*. One of these plasmids exhibited two replicons (IncHI2 and IncHI2A) and several genes that conferred resistance to different antibiotics: *tet*(M) (tetracycline); *dfrA12* (trimethoprim); *aadA2*, *aph(3’’)-Ib* and *aph(6)-Id* (aminoglycosides) and *floR* (phenicols).

**Figure 3.**
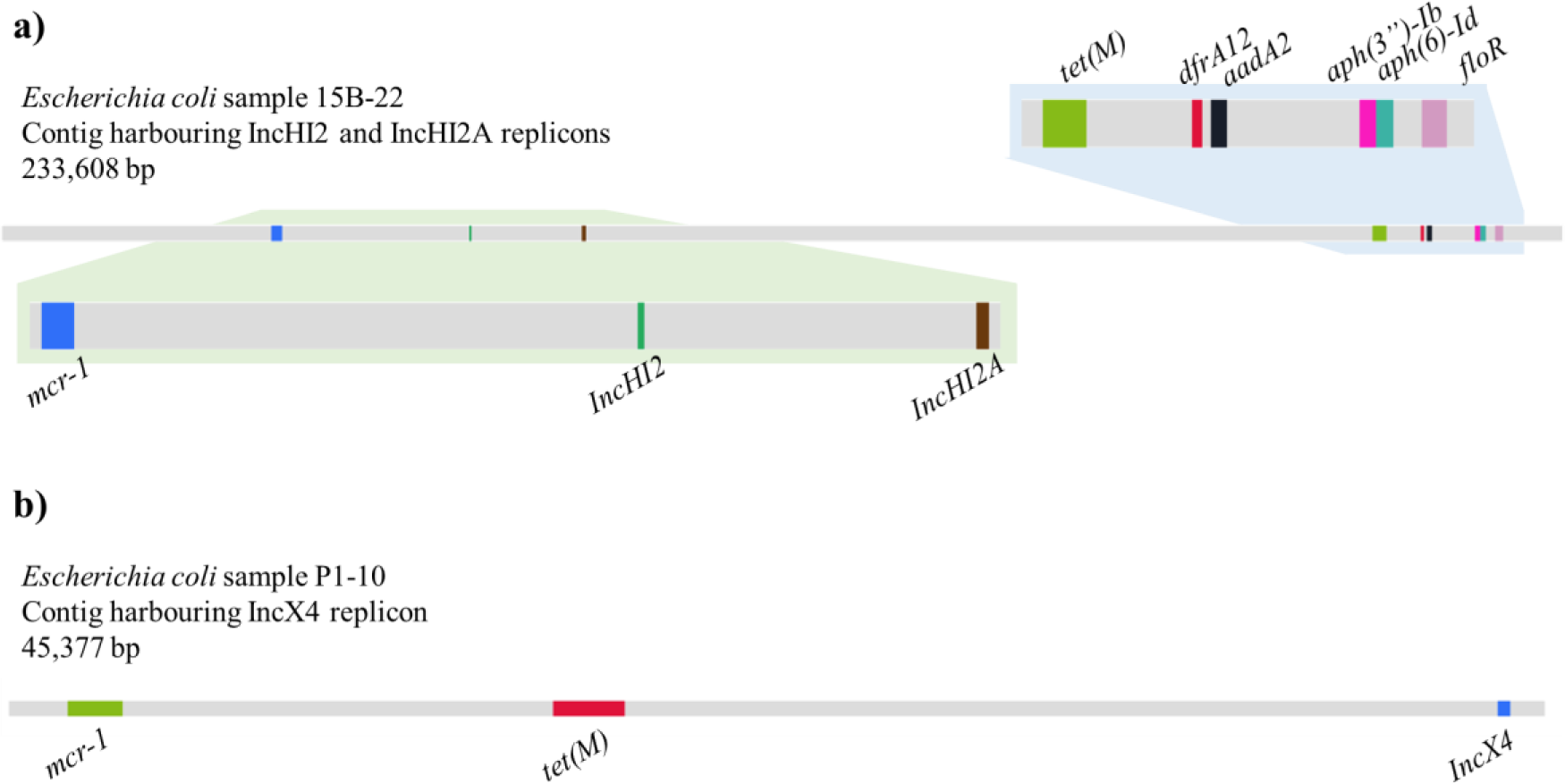
Contigs harbouring other resistances besides *mcr-1*. a) contig from isolate 15B-22 harbouring *mcr-1* gene, replicons IncHI2 and IncHI2A and several other genes that confer resistance to antibiotics: *tet(M)* (tetracycline); *dfrA12* (trimethoprim); *aadA2*, *aph(3’’)-Ib* and *aph(6)-Id* (aminoglycosides); *floR* (phenicols). b) contig from isolate P1-10 harbouring *mcr-1* gene, replicon IncX4 and *tet(M)* gene, which confers resistance to tetracycline.

Forty-four antimicrobial resistance genes were identified after sequencing the 18 colistin-resistant isolates (Table 2). For both replicons and antimicrobial resistance genes (Figure 4A and 4B respectively), the isolate from the farmer shares more elements with isolates from bovine rather than swine. In at least one of the representatives from human, swine and bovine we found the following resistance genes: against beta-lactams (*bla*_*TEM-1*_), aminoglycosides (*aad2, aph(3’)-Ia, aph(3’’)-Ib, aph(6)-Id*), trimethoprim (*dfrA12*), phenicols (*floR*), macrolides and lincosamides (*lnu(F)*), colistin (*mcr-1*) and sulphonamides (*sul3*). All antimicrobial resistance genes and replicons (but IncY) are detected in bovine isolates, representing an alarming reservoir for these elements.

**Figure 4.**
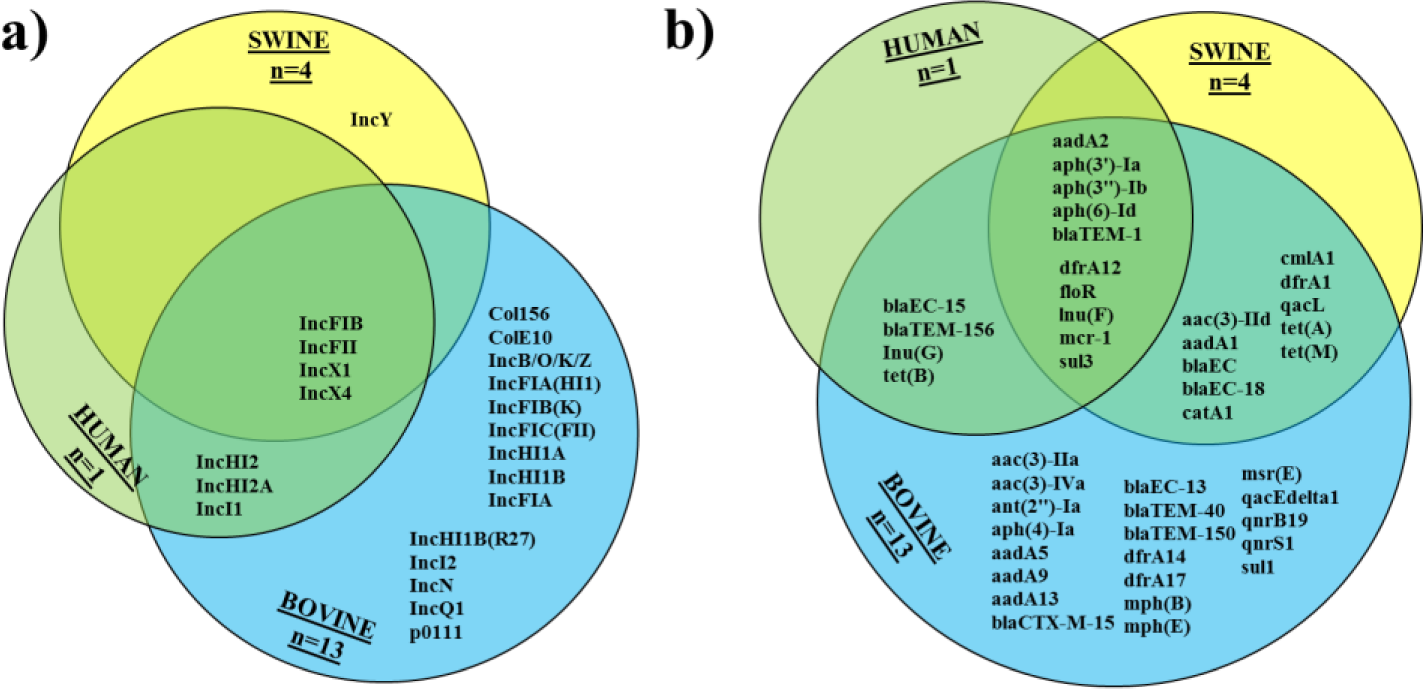
Venn diagrams for a) replicons and b) antimicrobial resistance genes identified by nanopore sequencing of the 18 colistin-resistant isolates.

## Discussion

The aim of this study was to conduct a thorough sampling of a farm to evaluate the co-occurrence of *mcr-1-mcr-3 E. coli* and assess the effectivity of the different molecular techniques including WGS to characterize *mcr*-positive isolates. Although occurrence of *mcr-3* could not be confirmed, *E. coli* harbouring *mcr*-1 was detected in different animal species co-inhabiting the facilities, including the farmer. According to the farm book, pigs and calves sampled in 2017 were both orally treated with colistin. Going back through the farm book, batches of both, pigs and calves reared from 2015 to 2017 were all prescribed colistin at the same time point of the production cycle. This management practice in terms of medication regime suggest a routine use of colistin in consecutive batches facilitating the emergence and persistence of colistin resistance mechanisms. Additionally, phenotypic and genotypic resistance to other families of antimicrobials such as beta-lactams, tetracycline, florfenicol, aminoglycosides and sulphonamides was very common. 15A-11 isolate, which resulted negative for *mcr-1* gene presence when performing sequencing and further analysis, was resistant to cephalosporins (cefotaxime and ceftazidime), harbouring *bla*_*CTX-M-15*_ gene.

The presence of *tet(M)* in the same plasmid harbouring *mcr-1* in P1-10 isolate could select for the presence of colistin resistance even if colistin is withdrawn, since doxycycline was also commonly prescribed to calves and pigs in this farm. In 15B-22 isolate, one of the two plasmids that encodes *mcr-1* gene (the one that harbours IncHI2 replicon), also encodes for another co-resistances: resistance to tetracycline (*tet(M)*), trimethoprim (*dfrA12*), aminoglycosides (*aph(3’’)-Ib*, *aph(6)-Id*) and florfenicol (*floR*). In that case, four families of antibiotics other than polymyxins (colistin) have a resistance gene within the same plasmid, meaning that there are more chances to maintain colistin resistance even if colistin is not used anymore.

Moreover, some strains present resistance to quaternary ammonium compounds (QACs) due to the presence of two genes (*qacEdelta1* and *qacL*, table 3). QACs are known because of its disinfectant and antiseptic proprieties^47,48^. It has been previously described the co-selection of antibiotic resistances to beta-lactams due to resistances to other kind of antimicrobials, such as heavy metals like silver, mercury, copper and zinc^49^. Copper and zinc are used as a mineral supplementation for animals, promoting the selection of multi-drug resistant bacteria. In that sense, and how is shown in Mulder et al.^48^, QACs are distributed in the environment and can end up accumulating in drinkable water and plants, leading to selection of these resistance strains once the animal drink or eat contaminated sources.

On the other hand, one of the isolates (15A-11) positive by PCR for *mcr-1* and phenotypically resistant to colistin was negative for the gene when nanopore sequencing was performed. Presumably, this isolate lost the plasmid during sub-culturing steps, which in the case of samples in hospital settings could result in false negative results and consequently in treatment failure.

Plasmids encountered herein contained similar replicons to the ones described in the literature to be potential carriers of colistin resistance, with IncX4 being the most prevalent^50–56^. In agreement with other studies^51,57–59^, some of the IncX4 plasmids have lost the transposition element ISApl1 involved in the mobilization of the *mcr-1* gene suggesting its stabilization within the plasmid. This was also the case for the isolate obtained from the farmer. Furthermore, a truncated fragment of the element remained in seven out of the 14 IncX4 plasmids, as a fingerprint of the whole structure that integrated in the plasmid once. In the case of IncI2 and IncHI2 the mobilization element was still present. Taking into account this information, IncX4 plasmids are the most evolved in terms of *mcr-1* stabilization due to the partial or full absence of the mobilization element that helps the gene to move among bacteria. Both IncI2 and IncHI2 plasmids present a full *ISApl1* element, meaning that the transposition event is likely to be more recent. Additionally, the bovine isolate 15B-22 yielded *mcr-1* in two different plasmids (IncHI2 and IncX4). Several studies have described coexistence of two *mcr-1-*harbouring plasmids within a host ^52–56^ or one copy on a plasmid and the other in the chromosome^53,55,56^.

As mentioned before, strains carrying *mcr-1* in the genome have also been described previously^50,53,55,56^. Interestingly, in our study, *mcr-1* was integrated in the genome in two isolates of different origin. In one occasion, the *ISApl1* element was flanking *mcr-1* upstream and downstream, structure that probably facilitates the movement of the whole element by transposition into other mobile genetic elements or along the genome. The extra isolate harbouring *mcr*-*1* in the chromosome had lost the *ISApl1* element and therefore has established *mcr-1* as a heritably trait that will be transferred to following generations. If the plasmid may suppose a fitness cost to the bacteria allowing its lost in the absence of selection^60^, harbouring colistin resistance in the chromosome may be an advantage overcoming this fitness cost. This isolate will become permanently resistant even when the selective pressure is removed ^50,61^.

Since different plasmid families and combinations of *mcr*-environments have been observed, it is most probable that *mcr*-positive strains have been introduced into the farm in different occasions. However, according to the ONT analysis of the *E. coli* genomes obtained in this study (data not shown), isolates bearing *mcr-1* in IncX4 plasmids belonged to different clones, suggesting repeated transfer of plasmids between them. Several studies have demonstrated transfer of resistant genes from livestock and poultry to farmers, veterinarians and personnel in direct contact with animals^49,62,63^ highlighting the impact of critically important resistance in public health and the importance of the One Health approach.

One-Health studies applying massive-parallel and single-molecule sequencing conform a good approximation to try to understand how antibiotic resistances flow between human-animal-environment scenario. There are different One-Health studies were Illumina^64^ and Nanopore^65^ or PacBio^66^ sequencing are used together to characterize bacteria isolated from different sources and try to unveil if there is relation amongst them. For example, in 2018, Ludden et al.^64^ performed a study in which concluded that there were similar antibiotic resistances between livestock and human *E. coli*, conferring resistance to 4 type of antibiotics, but on the other hand there was not a close relation between *E. coli* isolated from livestock and *E. coli* causing infection in human. Comparing their results with ours, eight out of 10 of the resistance genes shared between human, swine and bovine (Figure 4) are also present in their study, corresponding to resistance to aminoglycosides (*aph(3’)-Ia, aph(3’’)-Ib* or *strA, aph(6)-Id* or *strB*), beta-lactams (*bla*_*TEM-1*_), trimethoprim (*dfrA12*), florfenicol (*floR*), lincosamides (*lnu(F)*) and sulphonamides (*sul3*). This is translated to resistance among different species that share environment to 5 families of antibiotics, which is alarming due to the limiting effect on future infection treatment.

In 2018, Salinas *et al*.^66^ performed a study that included *E. coli* isolated from faecal samples from children and domestic animals. Even though the bacteria presented similar antibiotic resistance genes, they were genotypically diverse. For example, plasmids carrying the same antibiotic resistance genes were distinct among isolates from different origin.

In conclusion, *mcr-1* was found in different animal species in the farm including the farmer. Transfer of IncX4 plasmids from farm animals to the farmer is the most likely event, given that the animals were treated with colistin during the rearing cycle. ONT sequencing has proven to be a good and rapid tool for sequencing whole genomes and plasmids. The long reads retrieved facilitate the assembly of the DNA elements within the cell, and allows determining the presence of resistances within the same genetic element. Localization of antibiotic resistance genes and their genomic context is rapid and easy once the contigs are obtained after assembly.

## Data availability

The sequences of the 16 plasmids mentioned above and the Supplementary Data have been deposited in OSFHOME under the identifier <u>2F8W7</u>.

## Notes

https://osf.io/2f8w7/

